# Decomposing parietal memory reactivation to predict consequences of remembering

**DOI:** 10.1101/208678

**Authors:** Hongmi Lee, Rosalie Samide, Franziska R. Richter, Brice A. Kuhl

**Affiliations:** Department of Psychology, New York University; Department of Psychology, Boston College; Department of Psychology, Leiden University; Department of Psychology, University of Oregon

## Abstract

Memory retrieval can strengthen, but also distort memories. Parietal cortex is a candidate region involved in retrieval-induced memory changes as it reflects retrieval success and represents retrieved content. Here, we conducted an fMRI experiment to test whether different forms of parietal reactivation predict distinct consequences of retrieval. Subjects studied associations between words and pictures of faces, scenes, or objects, and then repeatedly retrieved half of the pictures, reporting the vividness of the retrieved pictures (‘retrieval practice’). On the following day, subjects completed a recognition memory test for individual pictures. Critically, the test included lures highly similar to studied pictures. Behaviorally, retrieval practice increased both hit and false alarm rates to similar lures, confirming a causal influence of retrieval on subsequent memory. Using pattern similarity analyses, we measured two different levels of reactivation during retrieval practice: generic ‘category-level’ reactivation and idiosyncratic ‘item-level’ reactivation. Vivid remembering during retrieval practice was associated with stronger category- and item-level reactivation in parietal cortex. However, these measures differentially predicted subsequent recognition memory performance: whereas higher category-level reactivation tended to predict false alarms to lures, item-level reactivation predicted correct rejections. These findings indicate that parietal reactivation can be decomposed to tease apart distinct consequences of memory retrieval.

## 1. INTRODUCTION

The act of retrieving a memory can affect the quality and accessibility of that memory in multiple ways. On the one hand, retrieved memories can become more resistant to forgetting (Karpicke and Roediger, 2008). On the other hand, retrieval can distort memory representations by creating an opportunity for new information to be incorporated into existing representations (Bridge and Paller, 2012; St. Jacques et al., 2013) or for some aspects of a memory to be strengthened more than others (Bouwmeester and Verkoeijen, 2011; Verkoeijen et al., 2012). For example, retrieval practice can promote false memory of semantically related but unstudied material if gist-level information is enhanced more than idiosyncratic details (Brainerd and Reyna, 1996; Payne et al., 1996; McDermott, 2006).

To understand the neural mechanisms underlying the consequences of retrieval, previous studies have examined the relationship between neural activity during retrieval practice and subsequent memory performance (van den Broek et al., 2016). In particular, retrieval-related activity in parietal cortex has emerged as one of the most reliable predictors of retrieval consequences (Wirebring et al., 2015). For instance, higher parietal activation during retrieval predicts better subsequent remembering (van den Broek et al., 2013; Liu et al., 2014). These findings align with the fact that successful retrieval activates parietal areas (Hutchinson et al., 2009; Rugg and King, 2017). In other words, retrieval-related responses in parietal cortex may predict memory strengthening because these responses reflect whether retrieval practice was successful. However, these findings do not tease apart the relative strengthening of gist-level versus idiosyncratic aspects of a memory, leaving open questions about the qualitative changes signaled by parietal activation.

Pattern-based analyses of memory representations have the potential to clarify the significance of parietal retrieval responses. In particular, accumulating evidence indicates that patterns of activity within parietal cortex reflect the contents of memory, and that reactivation of parietal activity patterns is closely related to the subjective experience of remembering (Kuhl and Chun, 2014; Chen et al., 2017). Notably, pattern-based measures of reactivation have generally come in two forms: (1) ‘category reactivation,’ which reflects information shared across stimuli from a common visual or semantic category (Polyn et al., 2005; Kuhl et al., 2011) and (2) ‘item reactivation,’ which reflects information idiosyncratic to specific stimuli or events (Kuhl and Chun, 2014; Wimber et al., 2015; St-Laurent et al., 2015). While these two measures of memory reactivation have often been used interchangeably, a few recent studies directly compared these measures and point to possible dissociations between them (Kuhl and Chun, 2014; Mack and Preston, 2016). To the extent that category and item reactivation are dissociable, an important possibility is that they predict distinct consequences of retrieval. Namely, whereas category reactivation putatively signals gist-level strengthening, item reactivation may signal strengthening of idiosyncratic information.

Here, we conducted an fMRI study to test whether category and item reactivation in parietal cortex predict distinct consequences of memory retrieval. Subjects first learned associations between words and pictures of faces, scenes, or objects. Subjects then repeatedly retrieved half of the pictures when cued with the associated words. A day later, subjects completed a recognition memory test for the previously-studied pictures. Critically, the recognition test included lures that were highly similar to studied pictures. Using pattern similarity analysis, we indexed the strength of category and item reactivation during retrieval practice within two regions of parietal cortex where category and item reactivation have previously been reported: angular gyrus and medial parietal cortex (Kuhl and Chun, 2014; Chen et al., 2017; Xiao et al., 2017). For comparison, we also measured reactivation within ventral temporal cortex–a set of high-level visual areas where category reactivation has frequently been observed (Kuhl et al., 2011; Schlichting and Preston, 2014). We hypothesized that stronger item reactivation in parietal cortex would reflect strengthening of idiosyncratic information and would guard against false recognition of lures, whereas category reactivation would reflect strengthening of gist-level representations and would therefore increase false recognition of lures.

## 2. MATERIALS AND METHODS

### 2.1. Participants

Twenty-eight healthy subjects (seven male, 18-32 years old) completed the study and were included in analyses. Two additional subjects were excluded: one due to excessive motion and one due to misunderstanding instructions. All subjects were right-handed and reported normal or corrected-to-normal vision. Informed consent was obtained in accordance with procedures approved by the University of Oregon Institutional Review Board.

### 2.2. Stimuli

We used a total of 288 words and 576 pictures. An additional 24 words and 24 pictures were used for practice trials. The words consisted of nouns, verbs, and adjectives between 3 and 11 letters in length *(M* = 6.14). The pictures consisted of color photographs collected from various online sources. There were three categories of pictures: famous people (e.g., Barack Obama; faces), famous places (e.g., the Great Pyramids of Giza; scenes), and common objects (e.g., scissors; objects). All of the pictures were of the same size (225 × 225 pixels). The pictures were grouped into two sets of 288 pictures, each consisting of an equal number (96) of faces, scenes, and objects. The two sets of pictures contained ‘corresponding’ pictures, such that for each picture in set 1, there was a similar (corresponding) picture in set 2. These corresponding pictures were not identical, but had a common referent. That is, corresponding pictures depicted the same person, place, or object (e.g., two different pictures of Barack Obama; see **Figure 1*C*** for examples). Each subject studied one of the two sets of pictures and the other set of pictures was used to generate the group of similar lures for the recognition memory test. Novel lures were also included in the recognition memory test and were drawn from the same set as the studied items. While the composition of the two sets was stable across subjects, the assignment of the sets to the studied list vs. similar lure list was counterbalanced across subjects. Individual words and pictures were randomly assigned to experimental conditions for each subject, with the constraint that the number of pictures from each visual category (faces, scenes, objects) was balanced across experimental conditions. Word-picture associations were also randomly determined for each subject. All stimuli were presented on a gray background. Text was presented in black. All visual stimuli were rear projected and viewed through a mirror attached to the head coil during scanning, and were presented on an iMac computer during behavior-only sessions.

**Figure 1.**
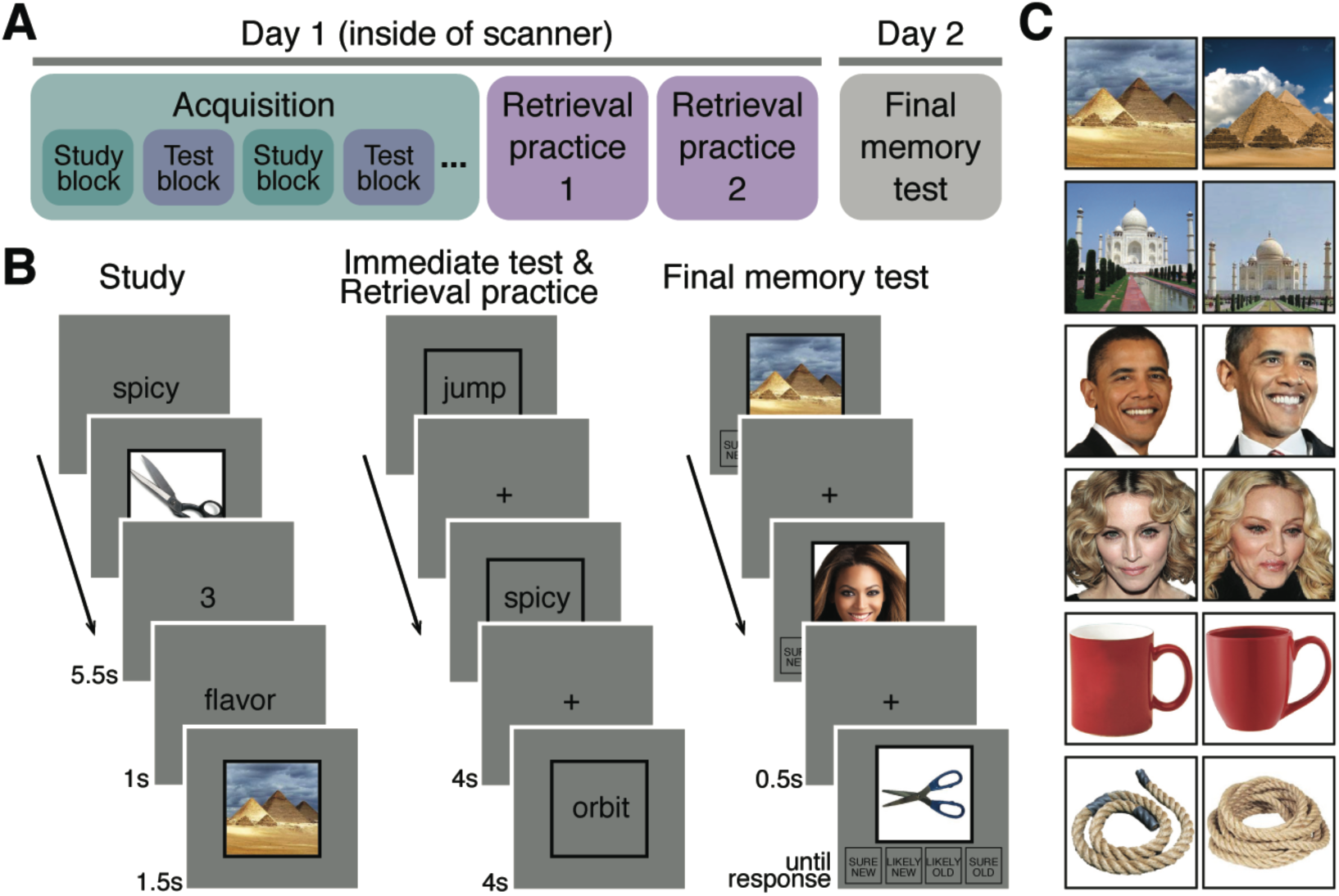
Procedures and example stimuli. (***A***) Overview of experimental phases. The acquisition phase (alternating study and immediate test blocks) and retrieval practice phases were conducted during fMRI scanning and performed on the same day. The final memory test was performed outside the scanner on the following day. (***B***) In the study blocks of the acquisition phase, subjects learned word-picture associations. In the immediate test blocks of the acquisition phase, and in the retrieval practice phase, subjects retrieved pictures associated with word cues and rated the vividness of their memories. In the final memory test phase, subjects made recognition memory judgments for pictures (without words). Pictures included previously-studied images (targets), highly similar lures, and novel images. (***C***) Example stimuli. The picture stimulus set consisted of pairs of highly similar, but not identical, pictures depicting the same person, place, or object. For each picture that was studied (during the acquisition phases), subjects were either tested with the same image (target) or the similar lure in the final memory test.

**Figure 2.**
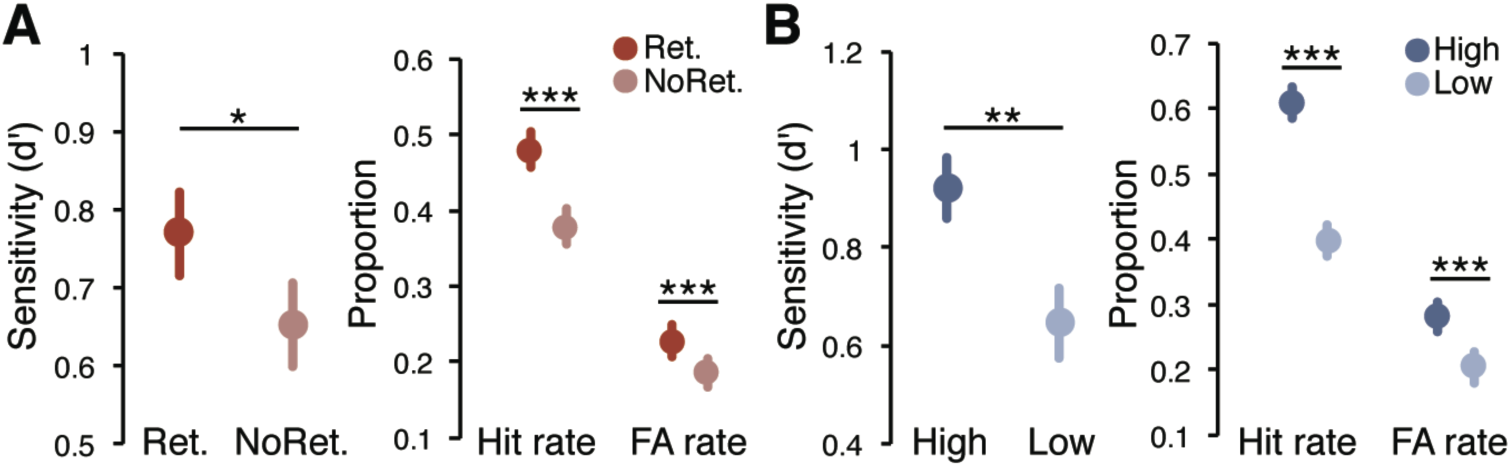
Behavioral results from the final memory test (picture recognition), combined across the behavioral pilot and fMRI studies. (***A***) Memory performance [average d’, hit rate, and false alarm (FA) rate] for pictures from the Retrieval vs. No retrieval conditions. (***B***) Memory performance (d’, hit rate, and FA rate) for pictures retrieved with High vs. Low vividness during RP1. The effects of vividness were qualitatively identical for RP2 trials (see Behavioral Results). Notes: In both ***A*** and ***B***, means and standard errors of the means (SEM) were computed from data collapsed across the behavioral pilot and fMRI experiments, but statistical significance was based on main effects of retrieval or vividness from mixed-design ANOVAs that included experiment (behavioral pilot vs. fMRI) as a between-subject factor. Error bars = SEM across subjects, \**p* < .05, \*\**p* < .01, \*\*\**p* < .001.

### 2.3. Experimental design and procedures

All experimental scripts were run in Matlab using the Psychophysics Toolbox (Brainard, 1997). The experiment spanned two consecutive days and consisted of three phases: acquisition, retrieval practice, and final memory test (**Figs. 1*A*-1*B***). The acquisition and retrieval practice phases were performed inside of the scanner on Day 1. The final memory test was performed on the following day and was not scanned.

#### 2.3.1. Acquisition

Subjects learned 192 word-picture associations through 16 cycles of ‘study’ and ‘immediate test’ blocks (8 scanning runs with 2 cycles each). Half of the studied pictures were retrieved (‘Retrieval’ condition) and the other half were not retrieved (‘No retrieval’ condition).

A study block consisted of 12 trials. Each trial started with a word cue (1s) followed by a picture (1.5s) presented in the center of the screen. After each picture, a 1-s fixation cross was presented, followed by an active baseline task. For the baseline task, two sequentially-presented numbers (randomly selected between 1 and 8) were presented for 1s each, with a 1-s gap between them. Subjects indicated whether each digit was an odd or an even number by pressing a button on an MRI-compatible response box with their right hand. An additional 1.5-s fixation cross followed the second digit. A total of 192 study trials (16 blocks × 12 trials/block) were presented in pseudo-random order with the constraint that each study block included an equal number of trials from each visual category condition. Additionally, within each study block, an equal number of trials were in the Retrieval vs. No retrieval conditions (see below) and an equal number of trials were subsequently tested with identical items vs. similar lures at the final memory test (see below). Each word-picture pair was only studied once.

In each immediate test block, subjects retrieved pictures associated with half of the word cues from the immediately preceding study block. Thus, each immediate test block consisted of six trials. The trial order was randomized within a block. Each trial started with a word cue presented for 4s at the center of a black square frame. The frame was of the same size as the picture stimuli. Subjects were instructed to retrieve the picture associated with the word cue as vividly as possible, and to rate the vividness of their memory on a 4-point scale (1 = least vivid, 4 = most vivid) via button press. There was no response option to specifically indicate retrieval failure. Responses were only recorded within 4s of the word cue onset. If no response was made within 3s, the color of the square frame changed from black to red for the last 1s to encourage subjects to respond within the given time window. A fixation cross was presented for 4s between trials, with no active baseline task. There were a total of 96 test trials.

Each study and immediate test block started with a 4-s instruction screen followed by a 6-s black fixation cross. All scanning runs ended with an additional 6-s fixation. At the end of each scanning run subjects were given feedback on the number of vividness responses that were made in the allotted 4-s window during the two immediate test blocks completed in that run. This feedback was intended to encourage vigilance.

All subjects completed a practice version of the acquisition phase in a prescreening session held on a separate day, prior to the main experiment. In the prescreening session, subjects completed two rounds of study and immediate test while inside a mock scanner. Each study and immediate test block had 12 and 6 trials, respectively. The inter-trial fixation during the practice session was shorter (1s) than that of the main experiment.

#### 2.3.2. Retrieval practice

The retrieval practice phase immediately followed the acquisition phase. During retrieval practice, subjects retrieved the same 96 pictures that were tested during the immediate test blocks. The trial structure was also identical to the immediate test trials from the acquisition phase. However, unlike the immediate test trials, retrieval practice trials were not interleaved with study trials. Thus, the lag between study and retrieval practice was substantially longer than the lag between study and immediate test. We anticipated that this delay would be a relevant factor in the relationship between retrieval/reactivation and subsequent memory, hence our use of different terms for the immediate test and retrieval practice phases. Subjects completed four retrieval practice scanning runs, each consisting of 48 trials. All 96 word cues from the Retrieval condition were presented once (RP1) within the first two runs and then all 96 word cues were presented again (RP2) in the last two runs. The presentation order of the word cues was randomized separately within the RP1 and RP2 lists. Each run started with a 4-s instruction screen followed by a 6-s fixation cross. A 6-s fixation cross was also added at the end of each run, followed by feedback on the number of responses that were made on time, as in the acquisition phase.

#### 2.3.3. Final memory test

On Day 2, subjects’ recognition memory for pictures was tested, but without presenting the word cues. Subjects completed a total of 288 trials. One-third of the test pictures were identical to the studied pictures (‘old’ items), one-third were similar lures that had the same referent as studied pictures without being identical (‘similar lures’), and one-third were novel pictures that were not presented on Day 1 and did not share a referent with any studied picture (‘novel’ items). Half of the pictures from the Retrieval condition were tested with old items and the other half were tested with similar lures; likewise for the No Retrieval condition. Visual category condition was also balanced across Retrieval/No Retrieval conditions and old/similar lure conditions. The final memory test phase was divided into 16 blocks, each consisting of 18 trials. The trial order was pseudo-randomized with the constraint that all experimental conditions were balanced within each block. No break or instruction was given between the blocks, so the block structure was not salient to subjects.

Each trial in the final memory test started with a picture presented at the center of the screen. Four response options were presented below the picture: ‘sure new,’ ‘likely new,’ ‘likely old,’ and ‘sure old.’ Each of the options was presented in a black square frame. Subjects selected one of the options via mouse click. Subjects were instructed to respond ‘old’ only when they had seen the exact same picture, and to respond ‘new’ to pictures that were similar to the studied pictures or entirely new. Thus, subjects were made aware of the similar lure condition prior to the final memory test. The task was self-paced. A 0.5-s inter-trial fixation cross replaced each picture after subjects made their response, but the four response options remained on the screen throughout the phase.

### 2.4. fMRI acquisition

fMRI scanning was conducted at the Robert and Beverly Lewis Center for NeuroImaging at University of Oregon on the Siemens Skyra 3T MRI scanner. Whole-brain functional images were collected using a T2*-weighted multi-band accelerated EPI sequence (TR = 2s; TE = 25ms; flip angle = 90°; 72 horizontal slices; grid size 104 × 104; voxel size 2 × 2 × 2 mm). A scanning session consisted of 8 acquisition runs and 4 retrieval practice runs. A total of 167 volumes were collected for each acquisition run and 200 volumes for each retrieval practice run. Additionally, a whole-brain high-resolution anatomical image was collected for each subject using a T1-weighted protocol (grid size 256 × 256; 176 sagittal slices; voxel size 1 × 1 × 1 mm).

### 2.5. fMRI analysis

#### 2.5.1. Preprocessing

Preprocessing of the functional data was conducted using FSL 5.0.9 (FMRIB Software Library, http://www.fmrib.ox.ac.uk/fsl) and custom scripts. Functional images were first corrected for head motion within each scanning run and then co-registered to the first scanning run. Motion-corrected images were smoothed with a Gaussian kernel (4mm full-width half-maximum) and high-pass filtered (cutoff = 0.01Hz). High-resolution anatomical images were brain extracted and co-registered to the functional images using linear transformation.

#### 2.5.2. General linear model analysis

Trial-specific fMRI activation patterns were estimated by running general linear model (GLM) analyses using each trial as a separate regressor. For each subject, separate GLMs were generated for the acquisition and retrieval practice phases using SPM12 (http://www.fil.ion.ucl.ac.uk/spm). The design matrix for each scanning run in the acquisition phase included 24 study and 12 immediate test trial regressors, and two additional regressors representing instructions for study and immediate test blocks, respectively. The design matrix for each scanning run in the retrieval practice phase included 48 retrieval trial regressors and an instruction regressor. All trial and instruction regressors were convolved with the canonical hemodynamic response function. For both GLMs, six motion parameters, impulse responses representing volumes with unusually large motion (i.e., motion outliers detected using the function fsl_motion_outliers in FSL), and constant regressors representing each scanning run were entered as regressors of no interest. Finally, one-sample *t*-tests were applied against a contrast value of zero to the resulting parameter estimates to obtain trial-specific *t* statistic maps. Univariate activation of each trial in a given region of interest was obtained by averaging the *t* values across all voxels within the region.

#### 2.5.3. ROI definition

We defined three a priori regions of interest (ROIs): angular gyrus (ANG), medial parietal cortex (MPC), and ventral temporal cortex (VTC). We first generated subject-specific anatomical ROIs via FreeSurfer cortical parcellation (http://surfer.nmr.mgh.harvard.edu). The ANG ROI consisted of bilateral angular gyrus as defined in FreeSurfer’s Destrieux atlas (Destrieux et al., 2010). The MPC ROI was a combination of bilateral precuneus, subparietal sulcus, and posterior cingulate gyrus. The VTC ROI was a combination of bilateral fusiform and parahippocamal gyri and adjacent areas including collateral sulcus and occipitotemporal sulcus. All anatomical ROIs were co-registered to the functional images. We further masked the ROIs with subject-specific whole-brain masks generated by SPM during GLM analyses to exclude voxels without *t* statistics. The number of voxels included in the anatomical ROIs varied across subjects (1,054–2,109 in ANG; 1,682–3,311 in MPC; 2,587–4,345 in VTC). For each subject, we selected the 500 most content-sensitive voxels within each anatomical ROI (**Fig. *3A***), which equated the number of voxels across ROIs and subjects. Content sensitivity of each voxel was determined using a one-way ANOVA on the *t* statistics of each voxel from the study trials using picture category as a factor. The 500 most content sensitive voxels were defined as the 500 voxels with the lowest *p* values. We additionally defined unilateral ANG ROIs by selecting the 250 most content-sensitive voxels separately from the left and right ANG to examine potential hemispheric differences in the effects of major experimental conditions.

**Figure 3.**
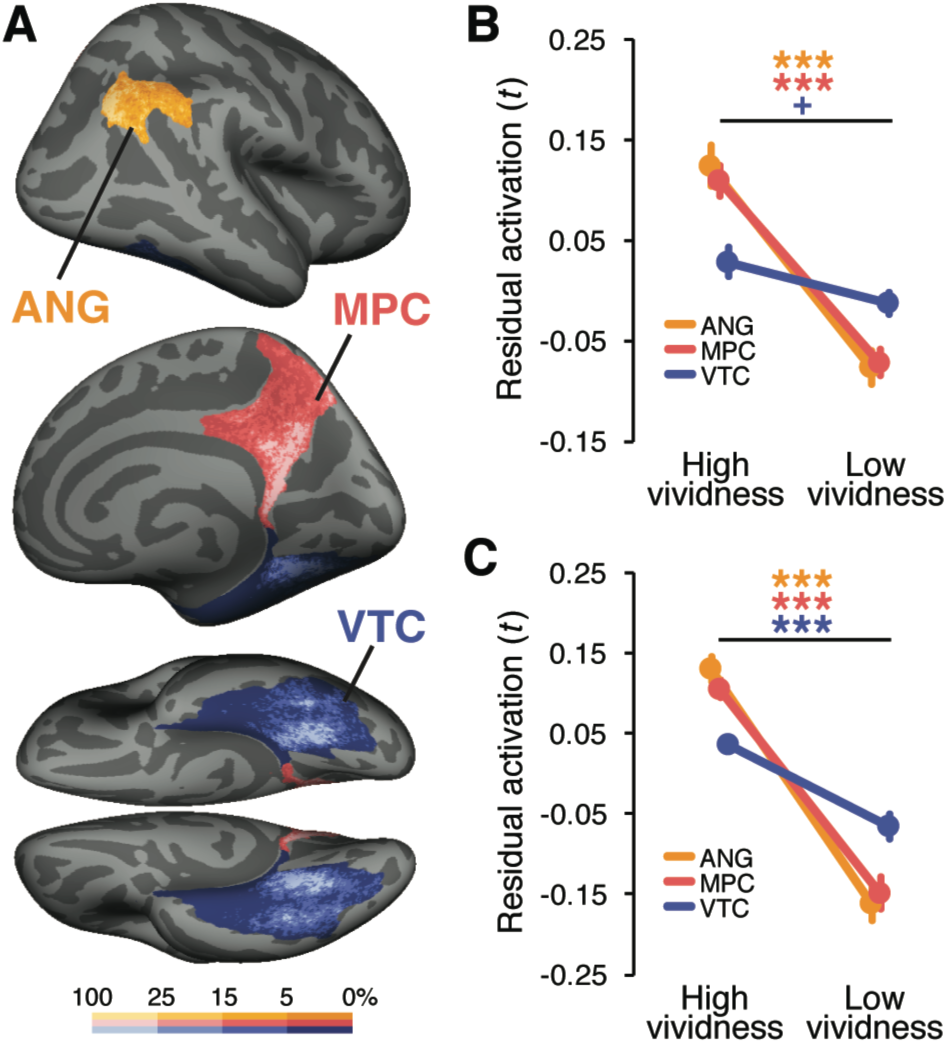
Regions of interest and univariate fMRI results. *(**A**)* Probability map of the 500 most category-sensitive voxels within each anatomical region of interest [orange = angular gyrus (ANG), red = medial parietal cortex (MPC), blue = ventral temporal cortex (VTC)], visualized on the inflated surface of the FreeSurfer template brain (fsaverage; top = lateral view of the right hemisphere, middle = medial view of the right hemisphere, bottom = ventral view). For each point on the surface of each region, the percentage of subjects whose category-sensitive voxels contained the point was computed. The color scale represents the percentages divided into four bins (smaller than 5%, 5-15%, 15-25%, greater than 25%), with brighter colors indicating higher percentages. (***B***) Univariate activation for High and Low vividness trials in the retrieval practice phase. (***C***) Univariate activation for High and Low vividness trials in the immediate test rounds of the acquisition phase. In both ***B*** and ***C***, univariate activation was defined as the residual activation after regressing out the main effect of picture category from the trial-specific *t* statistics averaged across voxels. Significance was determined from paired samples *t*-tests for each ROI. Error bars = SEM across subjects, +*p* < .1, \*\*\**p* < .001 (not corrected for multiple comparisons).

#### 2.5.4. Pattern similarity analysis

To measure the strength of category and item reactivation of the pictures during retrieval we used pattern similarity analyses (**Fig. *4A***). For each ROI, we first extracted the trial-specific *t* statistic patterns from three separate parts of the experiment: study trials, immediate test trials, and retrieval practice trials. Because the average vividness ratings and the relationship between vividness ratings and performance on the final test were comparable for RP1 and RP2 trials (see Behavioral results), we averaged activity patterns across corresponding RP1 and RP2 trials to create a single pattern per retrieved picture. Similarity between patterns was indexed using Pearson’s correlations. We separately considered similarity between study and immediate test trials and between study and retrieval practice trials. The resulting correlations were transformed to Fisher’s *z* before further analysis.

**Figure 4.**
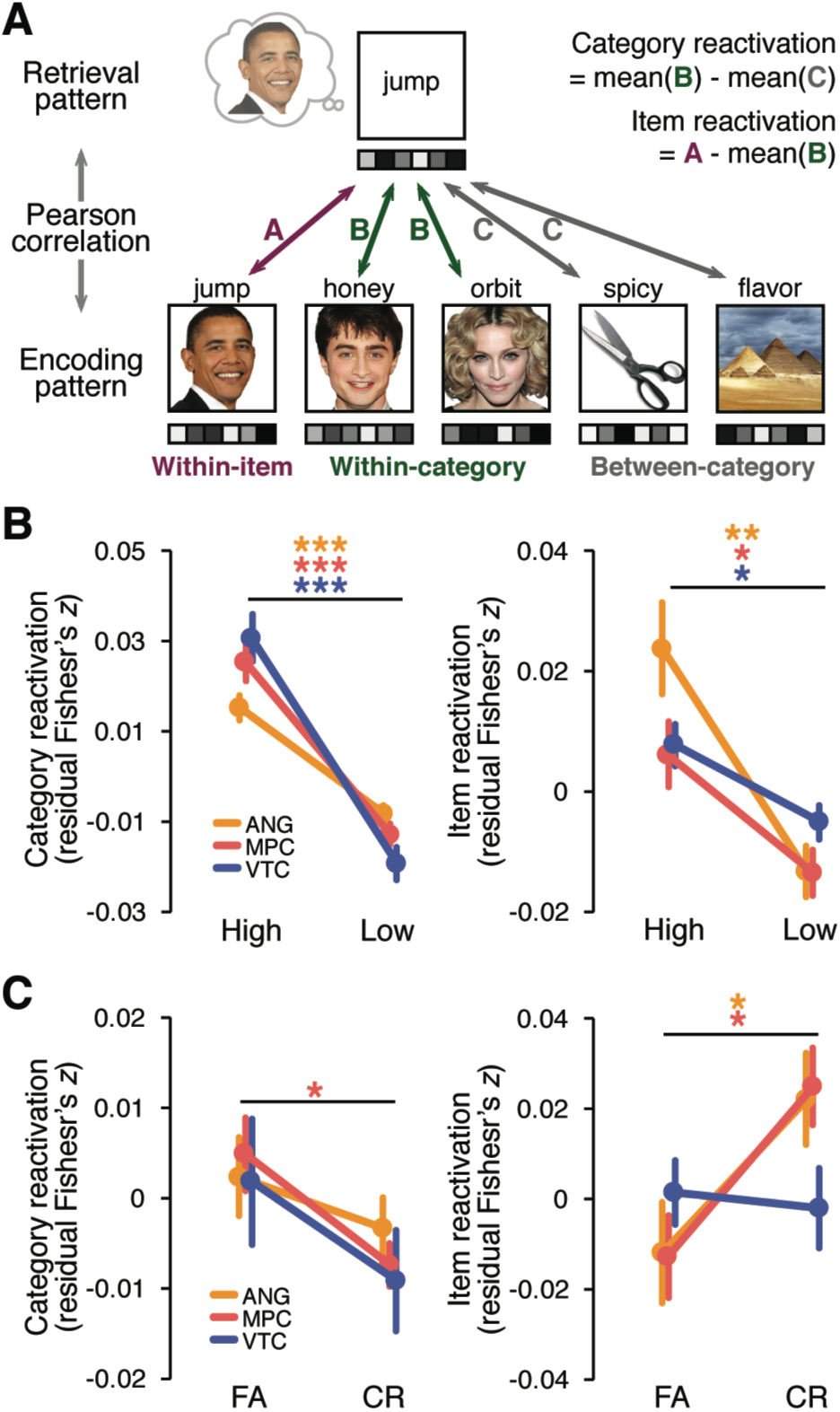
fMRI pattern similarity analyses. (***A***) Schematic of pattern similarity analyses. For each subject and ROI, the similarity (Pearson correlation) between each retrieval trial pattern and each study trial pattern was computed and transformed to Fisher’s *z*. For each retrieved picture, category reactivation was operationalized as the difference between its similarity to other pictures from the same category (within-category similarity) and its similarity to pictures from other categories (between-category similarity). Item reactivation was operationalized as the difference between within-item similarity and within-category similarity. (***B***) Category reactivation (left panel) and item reactivation (right panel) for High and Low vividness trials in the retrieval practice phase. (***C***) Category reactivation (left panel) and item reactivation (right panel) during retrieval practice as a function of subsequent false alarms (FA) and correct rejections (CR) during the final memory test. In both ***B*** and ***C***, category and item reactivation were expressed as residuals after regressing out main effects of picture category from each of the raw reactivation measures. Significance was determined from paired samples *t*-tests for each ROI (orange = ANG, red = MPC, blue = VTC). Error bars = SEM across subjects, \**p* < .05, \*\**p* < .01, \*\*\**p* < .001 (not corrected for multiple comparisons).

Category-level reactivation of each picture was operationalized as the difference between within-category and between-category similarity. Within-category similarity was computed by correlating the retrieval pattern of a given picture (from immediate test or RP1/RP2) with each study pattern corresponding to the same visual category as the retrieved picture; correlations were then averaged. The item-matched study trial was excluded from the within category similarity measure. Likewise, between-category similarity was computed by correlating the retrieval pattern for a given picture with each study pattern corresponding to pictures that were from a different visual category than that of the retrieved picture; correlations were then averaged. Similarly, item-level reactivation of a picture was defined as the difference between its within-item and within-category correlations, where within-item correlation was computed by correlating the study and retrieval patterns of the same picture. Thus category reactivation only reflected the degree to which generic visual category information was reactivated (Polyn et al., 2005; Kuhl et al., 2011) whereas item reactivation had the potential to capture item-specific or idiosyncratic information (Kuhl and Chun, 2014; Xiao et al., 2017). Importantly, the key results reported below were robust to changes in how category- and item-level reactivation were defined (Supplementary Results).

#### 2.5.5. Classification of subsequent memory outcomes

To test whether neural reactivation during retrieval practice predicted final memory test outcomes on a trial-by-trial basis, we trained a linear classifier to decode whether subjects would false alarm to or correct reject similar lure trials on the final memory test based on six measures of reactivation from the retrieval practice phase: category and item reactivation from each of the three ROIs (**Figs. *5A-5B***). Before running the classification analysis, we first regressed out measures of category and item reactivation that were collected during the immediate test rounds–this controlled for the effects of initial reactivation strength (see Results). Additionally, we regressed out the effect of visual category (face, object, scene). The residual category and item reactivation values were then z-scored across all trials within each subject (to remove across-subject differences). Each trial vector, which consisted of the six z-scored reactivation measures, was additionally normalized to a unit length. Classification was performed with an L-2 regularized logistic regression classifier (penalization parameter = 0.01) implemented in the Liblinear toolbox (Fan et al., 2008). We used a leave-one-subject-out cross-validation. That is, we trained the classifier on data concatenated across all but one subject (i.e., trials were pooled across subjects), and tested the classifier on each trial from the left-out subject. Classifier responses were considered correct when they matched the actual memory outcomes (false alarm or correct rejection). The percentage of correct classifier responses within each held-out subject was used as the decoding accuracy for that subject. To prevent classifier bias, we matched the number of false alarm and correct rejection trials within each training subject by randomly selecting the same number of items from each condition.

**Figure 5.**
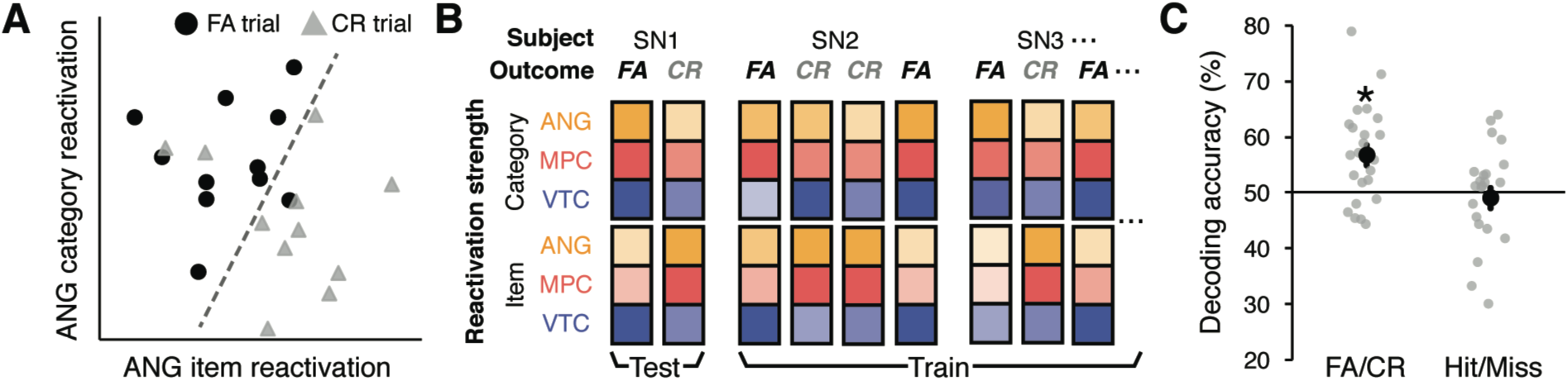
Across-subject decoding of subsequent memory outcomes. (***A***) Schematic of the across-subject classification approach. Scatterplot shows false alarm (FA) and correct rejection (CR) trials in a simplified 2-dimensional space where the *x* and *y* axes correspond to item reactivation in ANG and category reactivation in ANG, respectively. Each data point on the scatterplot represents a retrieval practice trial of a picture (averaged across repetitions). The data points were randomly selected from the actual data of an example subject such that the same number of FA (black circles) and CR trials (gray triangles) were included. The dotted line represents a hypothetical boundary that discriminates subsequent FA vs. CR trials based on the strength of item and category reactivation. For the actual analysis, 6 total features were used: category and item reactivation for each of the three ROIs (ANG, MPC, VTC). (***B***) Schematic of the leave-one-subject-out cross-validation procedure. For each iteration, the classifier was trained on the pooled data from all but one subject and tested on the left-out subject (in this example, SN1). Each training and test pattern consisted of six features: category and item reactivation strength from the three ROIs (shown as colored boxes). (***C***) Overall classification accuracy of subsequent FA vs. CR trials (left column) and transfer of the same classifier to Hit vs. Miss trials (right column). For the Hit vs. Miss decoding, FA and CR classifier ‘guesses’ were considered correct if they occurred on Hit and Miss trials, respectively. This analysis controlled for the classifier predicting subsequent responses (see Results). Black dots represent mean decoding accuracy (averaged across subjects, or iterations). Gray dots represent mean accuracy for each subject (averaged across multiple iterations, each with a random and balanced selection of FA and CR trials). Statistical significance reflects difference from chance, as determined from randomization tests. Error bars = SEM across subjects, \**p* < .05.

Because of this random sampling, we repeated the entire classification procedure 30 times (each with an independent random selection of the items) and averaged the decoding accuracy across repetitions to obtain a single accuracy per subject. Statistical significance of the classification performance was determined by a randomization test. We first ran a one-sample *t*-test against chance (50%) on the decoding accuracies across all subjects to obtain an ‘observed *t* value.’ We then generated a null distribution of *t* values by repeating the analysis 10,000 times using false alarm and correct rejection labels randomly shuffled within each training subject. The *p* value for this analyses was then defined as the proportion of *t* values of the null distribution that were equal to or greater than the observed *t* value from the un-shuffled data.

### 2.6. Behavioral pilot

Prior to conducting the fMRI experiment, we ran a behavioral pilot study highly similar to the fMRI experiment using an independent sample of subjects. The behavioral pilot was conducted in order to ensure that our retrieval practice manipulation had a causal influence on subsequent memory performance. Twenty-seven subjects with normal or corrected-to-normal vision participated (five male, 18-34 years old). Informed consent was obtained in accordance with procedures approved by the New York University Institutional Review Board. The experimental design, procedures, and stimuli were identical to those of the fMRI experiment except for the following: (a) the acquisition and retrieval practice phases were not divided into runs, (b) the presentation times for words and pictures during study trials (1.5s and 2.5s, respectively) were slightly longer than in the fMRI version, (c) the inter-trial fixation cross was shorter (1s) in the acquisition and retrieval practice phases, relative to the fMRI version, (d) the odd/even active baseline task that was used in the fMRI experiment was omitted, (e) no performance feedback was given, (f) response button labels were not presented on the screen between final memory test trials, (g) letters, fixation crosses, and frames were presented in white on a black background, (h) before the main experiment, subjects completed six encoding and three retrieval trials to practice the task.

### 2.7. Statistical analysis and data exclusion

In both the behavioral and fMRI data analyses, statistical comparisons between experimental conditions were performed with repeated-measures or mixed-design ANOVAs and two-tailed paired-samples *t*-tests. For all statistical tests, we report the raw *p* values. When post-hoc comparisons were performed for each ROI, we note if the effects of interest did not survive the Bonferroni correction.

For all fMRI analyses where univariate activation or pattern-based reactivation measures were related to subsequent memory or vividness, we regressed out effects of picture category and used the residuals as dependent variables. Importantly, this ensured that any observed relationships between the neural measures and subsequent memory or vividness would not be an artifact of differences across the picture categories. When the level of neural reactivation (category, item) was included as a factor in ANOVAs, we orthogonalized the category and item reactivation measures by additionally regressing out category reactivation from item reactivation within each subject. This procedure ensured that the two measures would not be correlated with each other despite that both measures were partially dependent on a same variable (i.e., within-category pattern similarity). However, virtually identical results were obtained when this additional orthogonalization step was omitted (see Supplementary Results).

Subjects who had fewer than five analyzable trials for any of the conditions of interest were excluded from the corresponding behavioral and fMRI analyses. Specifically, two subjects in the pilot experiment were excluded from the behavioral analysis comparing the high and low vividness conditions based on the retrieval practice phase ratings (see Behavioral results). One subject who had fewer than five high vividness trials was excluded from the fMRI analyses comparing the high and low vividness conditions in the retrieval practice phase. Five subjects who had fewer than five false alarm or correct rejection trials in the Retrieval condition were excluded from all fMRI subsequent memory analyses. In addition, for one fMRI subject, the first two trials of the RP2 phase were presented twice due to a technical issue. All study, immediate test, retrieval practice, and final memory test trials for the affected word-picture pairs were excluded from all behavioral and fMRI analyses (for that subject).

## 3. RESULTS

### 3.1. Behavioral results

Because behavioral memory performance was very similar across the behavioral pilot study and the fMRI experiment, we report the behavioral results combined across experiments (see **Table 1** for results separated by experiment). In analyzing final memory test performance, we only considered high confidence trials to reduce the influence of guess trials. Thus, hit trials were defined as ‘sure old’ responses for old pictures, miss trials were defined as ‘sure new’ responses for old pictures, false alarm (FA) trials were defined as ‘sure old’ responses for similar lures or novel pictures, and correct rejection (CR) trials were defined as ‘sure new’ responses for similar lures or novel pictures. The excluded low confidence trials constituted on average 43.4% *(SD* = 18.9%) of the trials per subject (sure old: 33.9%; likely old: 25.5%; likely new: 17.9%; sure new: 22.8%).

**Table 1.** Behavioral memory performance in the pilot and fMRI experiments.

|  |  | Retrieval conditions |  | RP1 vividness ratings |  | RP2 vividness ratings |  |
| --- | --- | --- | --- | --- | --- | --- | --- |
|  |  | Retrieval | No retrieval | High | Low | High | Low |
| Behavioral pilot | $d'$ | .79 (.37) | .62 (.41) | 1.00 (.44) | .62 (.50) | .97 (.50) | .68 (.46) |
|  | Hit rate | .51 (.19) | .41 (.18) | .64 (.15) | .44 (.19) | .64 (.17) | .42 (.21) |
|  | FA rate | .25 (.18) | .22 (.17) | .29 (.18) | .24 (.21) | .31 (.19) | .20 (.14) |
| fMRI experiment | $d'$ | .75 (.42) | .69 (.39) | .85 (.46) | .67 (.54) | .80 (.45) | .72 (.55) |
|  | Hit rate | .45 (.15) | .35 (.16) | .58 (.19) | .37 (.15) | .59 (.19) | .36 (.13) |
|  | FA rate | .20 (.12) | .16 (.10) | .27 (.15) | .17 (.12) | .30 (.15) | .16 (.12) |

Performance on the recognition memory test as a function of retrieval condition (left-most columns) and vividness ratings (middle and right-most columns) during retrieval practice. Mean $d'$ scores, hit rates, and false alarm (FA) rates are separately shown for the behavioral pilot study and the fMRI experiment. Standard deviation values are in parentheses.

To establish whether the similar lure pictures were effective in eliciting false alarms, we compared the false alarm rate for similar lure vs. novel picture trials. Indeed, the false alarm rate was markedly higher for similar lures than novel picture trials (*F*_1,53_ = 157.37, *p* < .0001). Subsequent analyses specifically focused on responses for old pictures vs. similar lures (i.e., excluding the novel picture trials). Overall, d’ values were significantly above 0 (*F*_1,53_ = 210.34, *p* < .0001), indicating above-chance discrimination of old vs. similar lure pictures. To critically test whether retrieval practice influenced final test performance, we compared final test performance across the Retrieval and No Retrieval conditions (**Fig. *2A***). Two-way mixed-design ANOVAs including the retrieval condition as a within-subject factor and the experiment as a between-subject factor revealed that retrieval practice significantly increased sensitivity as measured by d’ (Retrieval *M* = .77, No retrieval *M* = .65; *F*_1,53_ = 5.06, *p* = .029). Considered separately, retrieval practice increased both the hit rate for old items (Retrieval *M* = .48, No retrieval *M* = .38; *F*_1,53_ = 51.71, *p* < .0001) and the false alarm rate for similar lures (Retrieval *M* = .23, No retrieval *M* = .19; *F*_1,53_ = 14.4, *p* = .0004) with a significantly larger increase in the hit rate than false alarm rate (difference for hit rate, *M* = .10; difference for false alarm rate, *M* = .04; *F*_1,53_ = 13.88, *p* = .0005). No significant main effects of experiment nor interactions with experiment were found for any of these analyses (*p*s > .144).

We next assessed whether vividness ratings during the retrieval practice phase were predictive of final memory test performance (**Fig. 2*B***). To test this, we split RP1 and RP2 trials into high and low vividness groups (high = 3 and 4, low = 1 and 2), and conducted 2 (high, low vividness) × 2 (behavioral pilot, fMRI experiment) ANOVAs separately for RP1 and RP2. We found that for both RP1 and RP2 trials, higher vividness ratings were associated with higher d’ scores (*F*_1,51_s > 4.25, *p*s < .05), higher hit rates (*F*_1,51_ s > 70.36, *p*s < .0001), and higher false alarm rates (*F*_1,51_s > 18.38, *p*s < .0001), as revealed by significant main effects of vividness. Again, there was a greater effect of vividness on the hit rate than the false alarm rate (*F*_1,51_s > 12.61, *p*s < .001). No significant main effects of experiment or interactions by experiment were found for any of these analyses (*p*s > .12). The average vividness ratings and the consistency of vividness ratings across repeated retrieval trials are summarized in **Table 2**.

**Table 2.** Summary of vividness ratings.

|  | Mean vividness rating |  |  | % consistent response |  |  |
| --- | --- | --- | --- | --- | --- | --- |
|  | Immediate test | RP1 | RP2 | Immediate test & RP1 | Immediate test & RP2 | RP1 & RP2 |
| Behavioral pilot | 3.1 (.6) | 2.4 (.7) | 2.5 (.7) | 34.7 (14.4) | 35.4 (14.1) | 69.1 (10.9) |
| fMRI experiment | 2.7 (.5) | 2.2 (.4) | 2.2 (.4) | 52.2 (11.6) | 52.9 (14.4) | 71.6 (12.8) |

Left columns: mean vividness ratings made during the immediate test trials and the first and second retrieval practice trials (RP1 and RP2, respectively). Right columns: mean consistency of responses (i.e., percentage of corresponding trials for which subjects made identical vividness ratings) across the immediate test, RP1 and RP2 trials. Standard deviation values are in parentheses. Note: trials on which subjects did not make a response were excluded from these analyses. The percentage of 'no response' trials did not exceed 1.2 % in the immediate test, RP1, or RP2 trials.

Collectively, the behavioral data indicate that associative retrieval practice clearly had an influence on recognition memory for pictures a day later. Importantly, there were substantial increases not only in hit rates but also in false alarm rates, yet the increases in hit rates were relatively greater, resulting in modest improvements in sensitivity. This pattern of results suggests that there was strengthening of gist-level information as well as idiosyncratic information related to the retrieved pictures. It should be emphasized that if we had not included the similar lure trials–and instead only compared hit rates on old picture trials against false alarm rates on novel picture trials–the apparent benefit of retrieval would have been much stronger, but the relative strengthening of gist-level vs. idiosyncratic information would be much less clear. Thus, the similar lure condition represents a critical comparison point.

### 3.2. Univariate responses during retrieval

Univariate analyses were conducted to test whether the ROIs showed global increases in activation during vivid remembering (Kuhl and Chun, 2014). Specifically, we compared univariate activation (residual *t* values; see Methods) during high vividness retrieval practice trials (vividness ratings 3 and 4) vs. low vividness retrieval practice trials (vividness ratings 1 and 2) (**Fig. 3*B***; see **Supplementary Fig. 1*A*** for univariate activation separately for each of the four vividness ratings). For this analysis, we only included items that received consistent responses across RP1 and RP2 trials (high-high or low-low). This resulted in excluding on average 13.4% *(SD* = 7.2%) of items per subject. Consistent with prior findings (Kuhl and Chun, 2014), activation in parietal ROIs increased with vivid remembering: high vividness ratings were associated with significantly greater univariate activation in ANG and MPC (*t*_26_s > 6, *p*s < .0001). This effect was significantly greater in the left than right ANG (*F*_1,26_ = 37.44, *p* < .0001). The effect of vividness was marginally significant in VTC (*t*_26_ = 1.75, *p* = .092). The interaction between ROIs (ANG, MPC, and VTC) and vividness was significant (*F*_2,52_ = 26.83, *p* < .0001), reflecting the relatively greater effects of vividness in parietal regions. The same interaction between ROIs and vividness ratings was also observed in the immediate test rounds (*F*_2,54_ = 31.96, *p* < .0001; **Fig. 3*C***), although during immediate test all three ROIs showed higher activation for high vs. low vividness trials (*t*_27_s > 5, *p*s < .0001).

Although univariate activation was robustly related to subjective vividness ratings, the strength of univariate activation during retrieval practice did not predict subsequent performance for similar lures on the final memory test (i.e., correct rejections vs. false alarms). Specifically, a 2 (FA, CR) × 3 (ANG, MPC, and VTC) ANOVA on univariate activation did not reveal any significant main effects nor an interaction (*F*s < 1.5, *p*s > .25). Likewise, none of the individual ROIs exhibited a significant difference in univariate activation for FA vs. CR trials (*t*_22_s < 1). Thus, although activation in parietal cortex was highly sensitive to subjective vividness, overall levels of activation were not a reliable predictor of whether subjects would subsequently false alarm to or correctly reject similar lures.

### 3.3. Encoding-retrieval pattern similarity

We next examined whether information extracted from distributed activity patterns within the ROIs was predictive of whether subjects would false alarm to or correctly reject similar lures during the final memory test. Using pattern similarity analyses (comparing study trials to retrieval practice trials), we generated measures of two different levels of reactivation for each picture: category reactivation (i.e., within-category – between-category pattern similarity) and item reactivation (i.e., within-item – within-category pattern similarity). We first confirmed that both item and category reactivation were observed within the ROIs and also that these reactivation measures were modulated by retrieval vividness. To test whether item-level information was present in the ROIs during retrieval practice, we compared within-item vs. within-category similarity. A 2 (within-item, within-category) × 3 (ANG, MPC, and VTC) ANOVA revealed that within-item similarity was significantly higher than within-category similarity (*F*_1,27_ = 8.25, *p* = .008) with no interaction by ROI (*F*_2,54_ < 1), confirming item-level sensitivity. Similarly, category reactivation was evidenced by significantly greater within-category similarity than between-category similarity (*F*_1,27_ = 36.87, *p* < .001). An interaction with ROI (*F*_2,54_ = 8.7, *p* < .001) reflected relatively weaker category reactivation in ANG than in the other two regions.

To test whether the reactivation measures scaled with vividness during retrieval practice, we compared category and item reactivation during high vs. low vividness retrieval practice trials (**Fig. 4*B***). A 2 (category-level, item-level) × 2 (high vividness, low vividness) × 3 (ANG, MPC, and VTC) repeated-measures ANOVA revealed a highly significant main effect of vividness (*F*_1,26_ = 60.82, *p* < .0001) which did not interact with the level of reactivation (*F*_1,26_ = 2.4, *p* = .134) or ROI (*F <* 1). In all three ROIs, higher vividness ratings were associated with greater neural reactivation, as confirmed by significant main effects of vividness (*F*_1,26_s > 29.31, *p*s < .0001; no hemispheric difference in ANG, *F <* 1). There was also a significant reactivation level × vividness × ROI interaction (*F*_2,52_ = 7.85, *p* = .001), as the effect of vividness was more evident in category than item reactivation in VTC (*F*_1,26_ = 12.45, *p* = .002) but not in parietal ROIs (*F*_1,26_s < 2.09, *p*s > .16). **Supplementary Figures 1*B* and 1*C*** show category and item reactivation separated across each of the four vividness ratings.

Of central importance, we assessed whether category- and item-level reactivation of the pictures during retrieval practice differentially predicted subsequent false alarms to similar lures (**Fig. 4*C***). A 2 (category-level, item-level) × 2 (FA, CR) × 3 (ANG, MPC, and VTC) repeated-measures ANOVA revealed a significant interaction between the level of reactivation and subsequent memory (*F*_1,22_ = 7.80, *p* = .011)–that is, the two reactivation measures were related to subsequent false alarms in a qualitatively opposite manner. Specifically, category reactivation tended to be higher for subsequent FAs than CRs (*F*_1,22_ = 3.76, *p* = .066), whereas item reactivation was higher for subsequent CRs than FAs (*F*_1,22_ = 5.62, *p* = .027). There was a trend towards a 3-way interaction between reactivation level, subsequent memory, and ROI (*F*_2,44_ = 2.40, *p* = .102), with the interaction between reactivation level and subsequent memory only present in the parietal ROIs. Namely, considering the ROIs separately, the reactivation level × subsequent memory interaction was present in ANG (*F*_1,22_ = 6.22, *p* = .021; did not survive the Bonferroni correction; no hemispheric differences, *F <* 1) and MPC (*F*_1,22_ = 9.38, *p* = .006), but not in VTC (*F <* 1). The variability across ROIs was mainly driven by item reactivation, as higher item reactivation in ANG (*t*_22_ = 2.54, *p* = .019; did not survive the Bonferroni correction) and MPC (*t*_22_ = 2.68, *p* = .014) predicted CRs, while item reactivation in VTC did not differ between FAs and CRs (*t*_22_ < 1). Together, these results demonstrate that different levels of reactivation in parietal cortex predicted different memory outcomes: whereas coarser, category-level information tended to predict false alarms to lures, reactivation of item-specific information predicted resistance to false alarms.

### 3.4. Reactivation during immediate test rounds

For the above analyses, we specifically focused on reactivation during the retrieval practice trials, excluding reactivation during the immediate test rounds. There were two motivations for excluding reactivation from the immediate test rounds: (1) behavioral measures of vividness were markedly higher during the immediate test rounds than during retrieval practice (RP1 or RP2, *t*_27_s > 8, *p*s < .0001, see **Table 2**), suggesting that the retrieval practice trials presented more opportunity for memory modification, and (2) because the retrieval practice trials followed the immediate test trials, we anticipated that the immediate test trials would explain less variance on the final memory test than the retrieval practice trials. Here, we briefly consider reactivation during the immediate test rounds and its relation to memory performance. To first test whether category and item reactivation scaled with vividness ratings during the immediate test rounds, we ran ANOVAs with factors of reactivation level (category, item), vividness (high, low), and ROI (ANG, MPC, and VTC). As in the retrieval practice phase, there was a significant main effect of vividness (*F*_1,27_ = 48.74, *p* < .0001) which did not interact with reactivation level or ROI (*F*s < 1). When considering the ROIs separately, the main effect of vividness was significant in each ROI (*F*_1,27_s > 18.27, *p*s < .0005). The reactivation level × vividness × ROI interaction was also significant (*F*_2,54_ = 10.54, *p* = .0001), with the effect of vividness being greater in category than item reactivation in VTC (*F*_1,27_ = 10.85, *p* = .003) but not in parietal ROIs (*F*_1,27_s < 1.88, *p*s > .18). Thus, vividness effects in the immediate test rounds were qualitatively identical to those in the retrieval practice rounds.

When considering subsequent memory effects (FAs vs. CRs), a 2 (category-level, item-level) × 2 (FA, CR) × 3 (ANG, MPC, and VTC) ANOVA revealed a main effect of subsequent memory where reactivation was generally higher for subsequent FAs (*F*_1,22_ = 4.79, *p* = .04) without interacting with reactivation level (*F*_1,22_ = 1.19, *p* = .287). However, neither the main effect of subsequent memory (*F*_1,22_s < 3.86, *p*s > .062) nor the interaction between reactivation level and subsequent memory (*F*_1,22_s < 2.21, *p*s > .15) were significant in any of the individual ROIs. In short, the dissociation between category and item reactivation found in the retrieval practice phase was not present in the immediate test rounds. Rather–and consistent with our expectations–reactivation in the immediate test rounds was not a reliable predictor of performance on the final memory test.

The results above indicate that the strength of initial reactivation cannot explain the relationship between subsequent memory and reactivation during retrieval practice. That said, neural and behavioral responses during retrieval practice were not likely to be fully independent from those during immediate test. Indeed, vividness ratings were identical across the two phases for about half of the retrieved pictures (**Table 2**). In addition, within-subject, trial-by-trial correlations between the immediate test and retrieval practice phases were significantly positive for category and item reactivation in all ROIs (one-sample *t-*tests against 0; *t*_27_s > 3.59, *p*s < .002). Thus, to more strictly control for the potential influence of reactivation during immediate test on subsequent memory, we regressed out immediate test round reactivation from the reactivation in the retrieval practice phase on a trial-by-trial basis, separately for each subject and ROI. This resulted in category and item reactivation during retrieval practice that were orthogonal to category and item reactivation during the immediate test rounds, respectively. Using the residuals from these regressions, we obtained qualitatively identical results to our initial analyses. Specifically, there was a significant interaction between the level of reactivation and subsequent memory (*F*_1,22_ = 10.57, *p* = .004). Again, category reactivation tended to predict FAs (*F*_1,22_ = 3.7, *p* = .068) while item reactivation significantly predicted CRs (*F*_1,22_ = 8.39, *p* = .008), and this interaction tended to be stronger in parietal ROIs than in VTC, as reflected by a marginally-significant 3-way interaction between reactivation level, subsequent memory, and ROI (*F*_2,44_ = 2.67, *p* = .081). Considering the ROIs separately, the reactivation level × subsequent memory interaction was significant for ANG (*F*_1,22_ = 7.47, *p* = .012) and MPC (*F*_1,22_ = 10.57, *p* = .004), but not for VTC (*F*_1,22_ < 1).

These data, along with our behavioral findings, indicate that retrieval practice, in particular, influenced subsequent remembering and suggest that reactivation is a mechanism that underlies this influence. More specifically, our behavioral results indicate that retrieval practice causally increased the false alarm rate on the final memory test whereas our neural measures of reactivation indicate that reactivation within parietal regions explains unique variance in false alarm rates on the final memory test.

### 3.5. Cross-subject decoding of subsequent memory outcomes

The subsequent memory analyses described so far indicate that false alarms to similar lures on the final memory test are related to measures of neural reactivation during retrieval practice. As a stronger test of this idea, we next asked whether false alarms could be predicted on an item-by-item basis, and in a cross-validated manner, based on the measures of category and item reactivation during retrieval practice (**Fig. 5**). Specifically, we used a logistic regression classifier to predict FA vs. CR trials on the final memory test (using similar lure trials only) based on six measures of reactivation during retrieval practice: category and item reactivation scores for each of the three ROIs (ANG, MPC, VTC). Classification was performed using leave-one-subject-out cross-validation. To be clear, this classification analysis did not use patterns of fMRI activity; rather, this analysis only used ‘summary measures’ of reactivation strength that were extracted from patterns of fMRI activity via the analyses described above. Additionally, it is important to emphasize that the classifier was not being trained to discriminate FA vs. CR trials based on data from the final memory test (which was conducted outside the fMRI scanner and a day after retrieval practice); rather, the classifier was using retrieval practice measures to predict *subsequent* FAs vs. CRs. For the six reactivation measures, effects of picture category and strength of reactivation during the immediate test rounds were regressed out, with the residuals then used by the classifier. On average, 413.2 trials were used to train the classifier per iteration *(M =* 18.8, *SD* = 6.5 per subject). As with the above analyses, five subjects who had less than five FA or CR trials were excluded.

Mean decoding accuracy across subjects was significantly above chance *(M* = 56.78%, *p* = .019 from a randomization test), confirming that measures of reactivation during retrieval practice predicted, on a trial-by-trial basis, whether subjects would subsequently false alarm to or correctly reject individual similar lures. Notably, decoding accuracy was also above chance–and virtually identical–when only parietal ROIs were included in the analysis *(M* = 56.46%, *p* = .039). However, when only including category reactivation (for each of the three ROIs) or only item reactivation (for each of the three ROIs), classification accuracy was not reliably above chance *(M =* 52.72%, *p* = .126 and *M* = 53.60%, *p* = .117, respectively).

To test whether the classifier specifically learned to discriminate between subsequent FA and CR trials, we ran another classification analysis, this time training the classifier on FA vs. CR trials for n-1 subjects (just as before) but now testing the classifier on Hit vs. Miss trials for each held-out subject. Testing the classifier on Hit vs. Miss trials represents a useful control to rule out the possibility that the classifier simply learned to predict subsequent decisions (Old vs. New responses). Specifically, because FA trials and Hit trials each corresponded to trials where subjects responded ‘Old’ and because CR and Miss trials each corresponded to trials where subjects responded ‘New’, if the classifier simply learned to predict whether subjects would subsequently respond ‘Old’ vs. ‘New,’ then we would expect the classifier trained on FA vs. CR trials to perfectly transfer to Hit vs. Miss trials. Instead, when transferring from FA/CR to Hit/Miss trials, classification accuracy was not significantly different from chance *(M* = 49.10%, *p* = .351). Thus, the classifier’s ability to discriminate between subsequent FA vs. CR trials was not simply attributable to subsequent *responses*. Instead, it is more likely that category- and item-level reactivation reflected the relative strengthening of gist-level and idiosyncratic information, which was specifically relevant for determining how subjects would respond to similar lure trials. The results of these across-subject decoding analyses provide additional evidence that the observed relationship between reactivation level and subsequent memory was robust to different analysis approaches (also see Supplementary Results) while also providing novel evidence that trial-level false alarms can be *predicted* in held-out subjects based on the strength of category and item reactivation.

## 4. DISCUSSION

Here, we show that the act of remembering has multiple influences on memory representations and that these influences can be predicted by considering distinct forms of reactivation in parietal cortex. Specifically, our behavioral findings indicate that retrieval practice strengthened both gist-level and idiosyncratic representations, and our pattern-based fMRI measures of memory reactivation indicate that these behavioral consequences were predicted by generic category-level reactivation and item-specific reactivation, respectively. These findings build on a rich literature of behavioral investigations into the consequences of memory retrieval (Roediger and Butler, 2011), but provide new insight into the neural mechanisms that drive these consequences. Our findings add to accumulating evidence for memory reactivation within parietal cortex (Kuhl and Chun, 2014; Bird et al., 2015; St-Laurent et al., 2015; Chen et al., 2017), but our study is the first to consider whether parietal reactivation can be decomposed into distinct forms/measures that relate to distinct behavioral consequences. Below we consider the significance of our findings in relation to (1) the behavioral literature on consequences of retrieval practice, (2) pattern-based fMRI studies of memory reactivation, and (3) theoretical accounts of the role of parietal cortex in memory.

### Behavioral evidence for consequences of retrieval

Consistent with prior studies, our behavioral results indicate that retrieval practice significantly enhanced long-term retention (Karpicke and Roediger, 2008; Roediger and Butler, 2011), as evidenced by a higher hit rate for targets in the retrieval condition than in the no retrieval condition. This finding establishes the basic premise that retrieval practice causally influenced subsequent memory. However, our central focus was the influence that retrieval practice would have on memory for similar lures. Indeed, retrieval practice substantially increased the false alarm rate for similar lures (also see Roediger & McDermott 1995; McDermott 2006), suggesting that strengthening of gist-level information was a major component of the influence of retrieval practice (Bouwmeester & Verkoeijen, 2011; Verkoeijen et al., 2012). While strengthening of gist-level information would predict an increase in both hit rates and false alarms, it does not predict better discrimination of targets from similar lures given that these images shared a common conceptual label. Thus, the fact that we observed a modest increase in the ability to discriminate targets from similar lures, as measured by d’–or the disproportionate increase in hit rate compared to false alarm rate (**Fig. 2*A***)–suggests that strengthening also occurred for idiosyncratic information that discriminated targets from lures. Thus, the behavioral findings provide a strong hint that retrieval practice had separate–and potentially dissociable–influences on gist-level and idiosyncratic representations, motivating the critical question of whether these distinct influences are related to dissociable measures (or levels) of neural reactivation.

### Pattern-based fMRI measures of memory reactivation

Pattern-based fMRI analyses have enabled a major advance in studying the neural mechanisms of memory, as they provide a powerful means for measuring the contents and strength of memory reactivation (for a review, see Rissman and Wagner, 2012). In initial studies, reactivation was measured at the level of broad stimulus categories such as faces or scenes (Polyn et al., 2005; Kuhl et al., 2011). However, more recent studies have measured finer-grained reactivation at the level of individual events or items (e.g., Kuhl and Chun, 2014; Wing et al., 2015; Xiao et al., 2017). On the one hand, item reactivation may simply represent a more sensitive measure of reactivation and will therefore be more closely related to behavioral phenomena of interest (e.g., subsequent remembering). On the other hand–and consistent with our findings–item and category reactivation may relate to qualitatively distinct levels of memory representation and, therefore, track distinct behavioral phenomena (Koen and Rugg, 2016). While a few recent studies have found that the relative strength of category vs. item reactivation varies across brain regions (Kuhl and Chun, 2014; Mack and Preston, 2016), to our knowledge our findings constitute the first evidence that these different levels of reactivation are differentially related to the consequences of memory retrieval.

The dissociation we observed between item and category reactivation is particularly interesting given that both measures positively scaled with vividness ratings during retrieval practice. Additionally, univariate activity–which also scaled with vividness ratings and has widely been used as a marker of retrieval success (Buckner and Wheeler, 2001)–did not predict memory outcomes (FA vs. CR). Thus, our findings point to a specific relationship between the relative strength of item vs. category reactivation and the ability to subsequently discriminate retrieved items from highly similar lures. This relationship was robust to differences in how item and category reactivation were measured (see Supplementary Results) and our cross-subject decoding results demonstrate that by simply considering the strength of item vs. category reactivation, behavioral outcomes (FA vs. CR) can be predicted on individual trials–from ‘held out’ subjects–with above-chance accuracy.

One caveat to our findings is that the neural measures of item and category reactivation described here are not at precisely the same representational level as the idiosyncratic vs. gist information probed by the final memory test. Namely, item reactivation was defined as identity-specific or location-specific reactivation, but correctly rejecting similar lures on the final memory test required even finer-grained discrimination between images that shared identities or locations. Likewise, category reactivation was defined as reactivation that differentiated between the two broad classes of stimuli (faces vs. scenes) whereas gist-based false alarms reflected a failure to discriminate between images within a category that shared an identity or location. Thus, our argument is only that item and category reactivation differed in their relative sensitivity to idiosyncratic vs. gist-level information, as opposed to these being ‘pure’ measures of idiosyncratic vs. gist-level information. Additionally, the comparison of category vs. item reactivation is useful given that these measures have been widely used in other pattern-based fMRI studies on memory reactivation (see above). However, future work may explore memory reactivation across a wider range of granularities.

### Relevance to theoretical accounts of the role of parietal cortex in memory

Pattern-based fMRI studies have revealed that parietal cortex contains surprisingly rich information about retrieved content (Kuhl and Chun, 2014; Lee and Kuhl, 2016; Bonnici et al., 2016; Thakral et al., 2017), but there remains debate about the nature of these representations. One possibility is that parietal cortex represents abstract semantic concepts of remembered events. This account is motivated by evidence that parietal cortex–and ANG, in particular–is a component of the semantic memory network (Binder and Desai, 2011). Indeed, activation patterns in parietal cortex reflect the semantic categories of stimuli in a modality-general manner (Devereux et al. 2013). However, prior studies have also found that parietal cortex is involved in processing of perceptual details. For example, inferior parietal activation during retrieval is greater for true recognition than gist-based false recognition (Guerin et al. 2012). ANG activation also scales with the precision of retrieved memories, as measured in terms of visual features such as color, location, or orientation (Richter et al. 2016).

However, findings from our study are equivocal with respect to whether parietal representations are semantic vs. perceptual. While the fact that category reactivation in parietal cortex tended to be associated with gist-based false alarms suggests a semantic or conceptual representation of retrieved memories, the fact that item reactivation in parietal cortex predicted discrimination of ‘semantically equivalent’ images suggests at least some sensitivity to perceptual details. While it is tempting to interpret this dissociation as evidence that category reactivation maps to semantic reactivation whereas item reactivation maps to perceptual reactivation, this is likely an oversimplification. Indeed, it is more likely that each measure includes a combination of semantic and perceptual information, though perhaps to varying degrees. More generally, our findings are consistent with the idea that parietal cortex combines multiple forms of remembered information (Shimamura, 2011; Wagner et al., 2015; Bonnici et al., 2016). In fact, the combination of diverse inputs may be precisely why parietal activity patterns are ‘content rich.’

Finally, a critical aspect of the parietal results is that they were not mirrored by VTC. In particular, item reactivation in VTC did not predict correct rejections of similar lures. The selectivity of these effects to parietal regions was consistent across different methods for measuring item and category reactivation (see Supplementary Results). This dissociation between parietal cortex and VTC is notable given that memory reactivation has more frequently been measured in VTC (Wheeler et al., 2000; Wimber et al., 2015), motived by the idea that remembering involves reinstating sensory experience (Danker and Anderson, 2010). However, our findings–and findings from a few prior studies (Kuhl et al., 2013; Kuhl and Chun, 2014; Bird et al., 2015)–suggest that parietal memory reactivation may be more closely related to behavioral performance than reactivation in VTC. Notably, even among frontal and temporal regions functionally connected to ANG and MPC (i.e., sub-regions of the default mode network; Yeo et al. 2011), we did not observe interactions between reactivation level (item vs. category) and subsequent memory (**Supplementary Figure 2**). Indeed, when considering the relationship between reactivation level and subsequent memory, there was a significant difference between parietal vs. frontal/temporal regions of the default mode network (**Supplementary Figure 2**), even though frontal/temporal regions demonstrated univariate and pattern-based effects of vividness similar to VTC and parietal regions. Thus, the observed pattern of results in parietal cortex was not a general property of the default mode network nor was it mirrored by VTC. Rather, although reactivation is distributed across many brain regions, reactivation in parietal cortex may be particularly related to the consequences of retrieval.

One possibility–though speculative–is that parietal cortex may function as an interface between sensory/perceptual reactivation (Wheeler et al., 2000) and top-down goal signals from prefrontal cortex (Tomita et al., 1999). Indeed, parietal representations are flexible across changing tasks (Ibos and Freedman, 2014) and are more sensitive to retrieval goals than VTC (Kuhl et al., 2013). An important avenue for future research is to understand how parietal cortex interacts with other brain areas to represent retrieved memories.

## Acknowledgements

This work was made possible through generous support from the Lewis Family Endowment to the UO which supports the Robert and Beverly Lewis Center for Neuroimaging.

## SUPPLEMENTARY RESULTS

### Pattern similarity results based on alternative measures of reactivation

For all analyses reported in the main text, we defined category reactivation as [within category similarity – between category similarity] and item reactivation as [within item similarity – within category similarity]. However, these definitions create a bias toward a negative relationship between the two measures. Indeed, there was a small, but significant negative correlation between these measures (after controlling for the effect of stimulus category) in ANG [mean correlation (Fisher’s *z*) = -.06, *t*_27_ = 3.6, *p* = .005, one sample *t*-test against 0] and MPC [mean correlation (Fisher’s *z*) = -.05, *t*_27_ = 2.92, *p* = .007], but not VTC [mean correlation (Fisher’s *z*) = .01, *t*_27_ = .38, *p* = .71]. Because of the non-independence between these measures, for all ANOVAs reported in the main text that included reactivation level as a factor (category vs. item) we regressed category reactivation out of item reactivation resulting in orthogonal reactivation measures. However, to confirm that our critical results were robust to the specific way in which reactivation levels were measured, we repeated key analyses using different measures.

First, omitting the step of regressing category reactivation out from item reactivation had very little influence on the interaction between reactivation level and subsequent memory. Namely, with this step omitted a 2 (category-level, item-level) × 2 (FA, CR) × 3 (ANG, MPC, and VTC) repeated-measures ANOVA still revealed a significant interaction between reactivation level and subsequent memory (*F*_1,22_ = 8.11, *p* = .009) and a marginally significant 3-way interaction (*F*_2,44_ = 2.62, *p* = .084). Considering individual ROIs, the interaction between reactivation level and subsequent memory was observed in ANG (*F*_1,22_ = 5.64, *p* = .027; not significant after Bonferroni correction) and MPC (*F*_1,22_ = 9.96, *p* = .005), with category reactivation numerically higher for FAs than CRs and item reactivation higher for CRs than FAs. In contrast, there was no interaction in VTC (*F <* 1) where both reactivation measures were numerically higher for FAs than CRs.

Second, we also re-ran these same analyses using a different definition of item and category reactivation. Namely, item reactivation was defined as within-item similarity with no subtraction of within-category similarity, and category reactivation was defined as within-category similarity with no subtraction of between-category similarity. Using these alternative measures, the bias toward a negative relationship between the reactivation measures was eliminated. Indeed, with these alternative definitions, there was now a positive correlation between category and item reactivation in ANG, MPC, and VTC (*t*_27_s > 9.25, *p*s < .0001). However, a 2 (category-level, item-level) × 2 (FA, CR) × 3 (ANG, MPC, and VTC) repeated-measures ANOVA again revealed an interaction between reactivation level and subsequent memory (*F*_1,22_ = 5.62, *p* = .027) as well as a 3-way interaction (*F*_2,44_ = 4.82, *p* = .013). Considering individual ROIs, the interaction between reactivation level and subsequent memory was present in ANG (*F*_1,22_ = 6.46, *p* = .019; not significant after the Bonferroni correction) and MPC (*F*_1,22_ = 7.17, *p* = .014), but not in VTC (*F <* 1). For ANG and MPC, the interaction was qualitatively identical to the interaction reported in the main text (and in the analysis above), with both regions characterized by relatively stronger category reactivation for subsequent FAs than CRs and relatively stronger item reactivation for subsequent CRs than FAs. Thus, regardless of the specific way in which reactivation measures were computed, we consistently observed the same critical interaction between reactivation level and subsequent memory. Namely, whereas item-level reactivation predicted subsequent correct rejections of lures, category-level information tended to predict false alarms. Moreover, this pattern of results was consistently present within parietal ROIs, in particular.

## SUPPLEMENTARY FIGURES

**Supplementary Figure 1.**
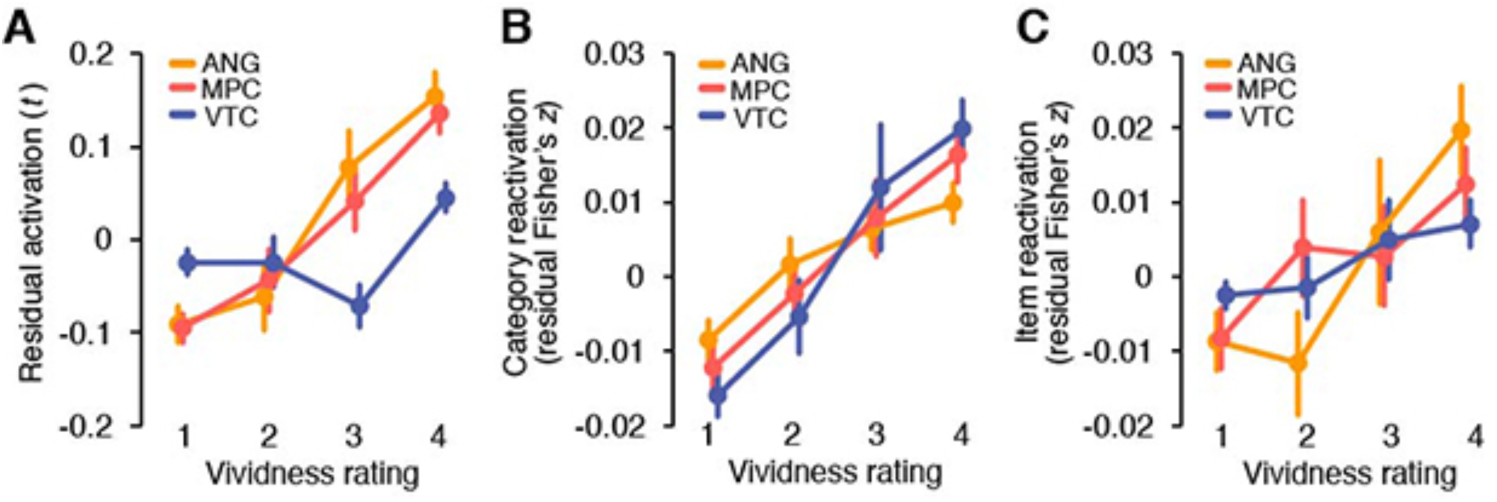
Relationship between vividness ratings during retrieval practice trials and (***A***) Univariate activation, (***B***) Category reactivation, and (***C***) Item reactivation. Vivid ratings range from 1 (least visit) to 4 (most vivid). Results were based on the data concatenated across R1 and R2. All dependent variables were expressed as residuals after regressing out main effects of picture category from each of the raw variables. Repeated-measures ANOVAs with factors of vividness rating (1 to 4) and region (ANG, MPC, and VTC) revealed significant main effects of vividness for each of the three dependent variables (*F*_3,72_s > 4.24, *p*s < .009) as well as significant positive linear trends (*F*_1,24_s > 20.4, *p*s < .001) but not quadratic trends (*F*_1,24_s < 1.94, *p*s > .17). A significant vividness × ROI interaction was observed for univariate activation (*F*_3.3,78.6_ = 9.62, *p* < .0001; Greenhouse-Geisser correction applied), but not for category reactivation (*F*_3.8,90.6_ = 1.41, *p* = .213; Greenhouse-Geisser correction applied) or item reactivation (*F*_6,144_ = 1.20, *p* = .309). For univariate activation the vividness × ROI interaction was driven by differences between parietal ROIs and VTC, with significant positive linear trends in both parietal ROIs (*F*_1,24_s > 37.4, *p*s < .0001) but not in VTC (*F*_1,24_s = 2.85, *p* = .105). Notably, for all three dependent measures, the main effects of vividness and the positive linear trends remained significant when only vividness ratings 2 to 4 were considered (*F*_2,48_s > 3.21, *p*s < .049). Thus, the effects of vividness can not be explained by a difference between ratings of “1” (which may indicate complete retrieval failure) vs. ratings of “2”, “3” or “4.” When only considering ratings of 2 to 4, the effect of vividness again only interacted with ROI when considering univariate activation (*F*_2.8,68_ = 7.72, *p* = .0002; Greenhouse-Geisser correction applied), with positive linear trends in both parietal ROIs (*F*_1,24_s > 15.4, *p*s < .001) but not in VTC (*F*_1,24_ = 3.52, *p* = .073). Error bars = SEM across subjects (*N* = 25 after removing three subjects who had less than five trials in any of the four vividness levels).

**Supplementary Figure 2.**
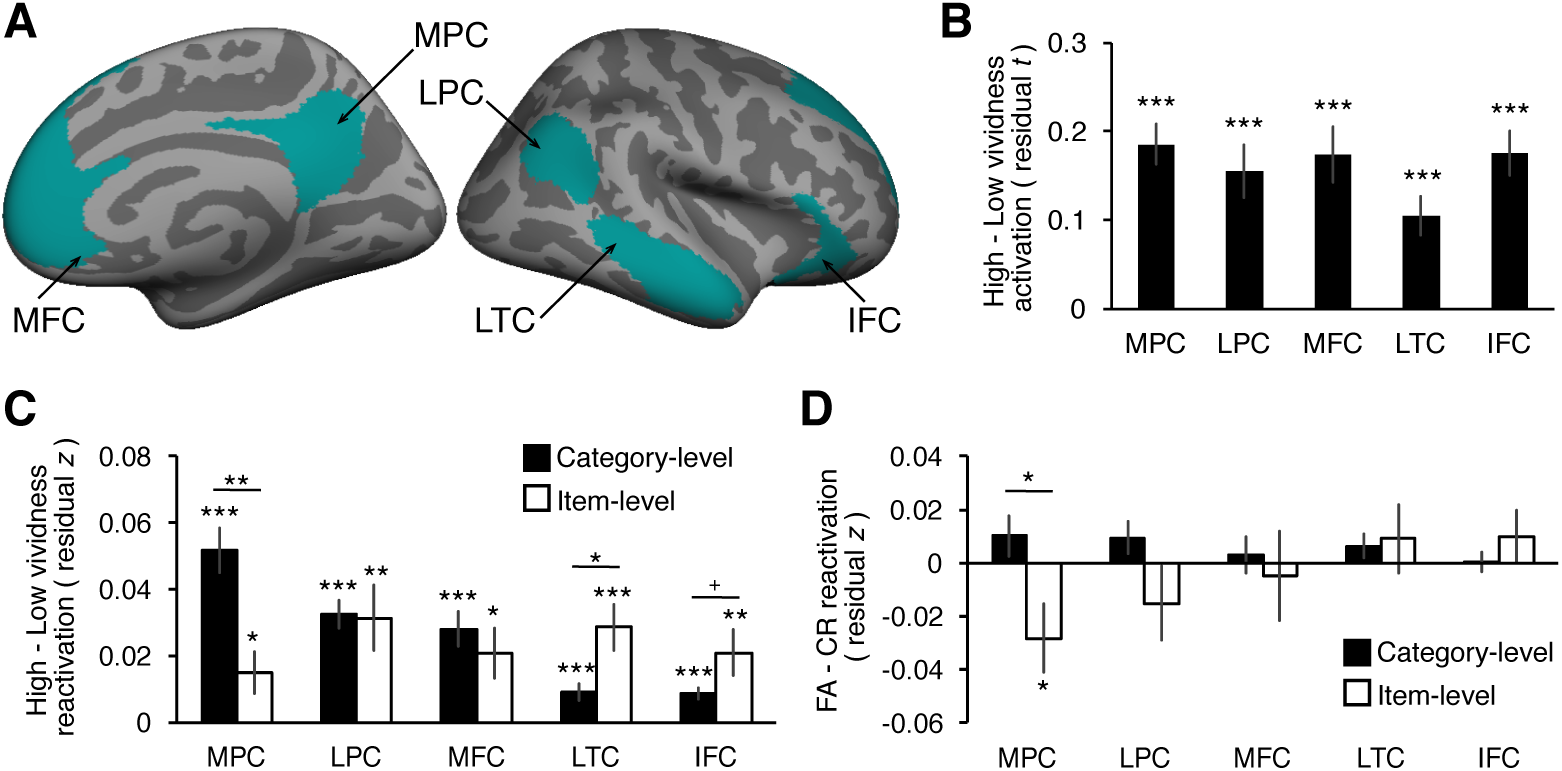
Effects of vividness and subsequent memory in sub-regions of the default mode network. (***A***) Regions of interest [medial parietal cortex (MPC), lateral parietal cortex (LPC), medial frontal cortex (MFC), lateral temporal cortex (LTC), interior frontal cortex (IFC)] were selected from the default mode network as defined by intrinsic functional connectivity data (Yeo et al, 2011; seven-network version), visualized on the inflated surface of the FreeSurfer template brain (fsaverage; left = medial view of the right hemisphere, right = lateral view of the right hemisphere). The ROIs defined on the surface of each hemisphere of the template brain were first co-registered to the surface of each subject’s anatomical image and then registered to the 3D functional image space. The resulting unilateral ROIs were masked by subject-specific whole-brain masks generated by SPM and combined across hemispheres. Within each bilateral ROI mask, the 500 most category-sensitive voxels were selected using a one-way ANOVA on the univariate activation of each voxel from the study trials. (***B***) Vividness effects (the difference between High vs. Low vividness ratings) on univariate activation during retrieval practice. Each region showed higher activation during high than low vividness trials (*t*_26_s > 4.7, *p*s < .001). (***C***) Vividness effects (the difference between High vs. Low vividness ratings) on category and item reactivation during retrieval practice. Both reactivation measures were higher for high than low vividness trials in all regions (*t*_26_s > 2.77, *p*s < .011; effects on item reactivation in MPC and MFC did not survive Bonferroni correction). (***D***) Subsequent memory effects (the difference between FA vs. CR trials) on category and item reactivation during retrieval practice. When the two parietal regions (MPC and LPC) were considered together, there was a significant interaction between reactivation level and subsequent memory (*F*_1,22_ = 5.22, *p* = .032) with no interaction between the regions (*F <* 1) [note: the interaction between reactivation level and subsequent memory in the parietal regions was not significant after Bonferroni correction for the number of ROIs]. No interaction between reactivation level and subsequent memory was observed when frontal and temporal regions were considered together or separately (*F*s < 1). This dissociation between parietal and frontal/temporal regions was confirmed by a significant 3-way interaction between region (parietal, frontal/temporal), reactivation level, and subsequent memory (*F*_1,22_ = 6.4, *p* = .019) [note: for this analysis, data was averaged across sub-regions separately for parietal (MPC, LPC) and frontal/temporal (MFC, LTC, IFC) regions]. In ***B*** – ***D***, univariate activation and reactivation measures were expressed as residuals after regressing out main effects of picture category from each of the raw dependent variables. Significance against 0 (i.e., no difference between conditions) was determined from one-sample *t*-tests for each ROI and dependent variable. In ***C*** and ***D***, differences between category and item reactivation for each ROI were determined from 2 (category-level, item-level) × 2 (high/low vividness or FA/CR) repeated-measures ANOVAs after orthogonalizing category and item reactivation by regressing out category reactivation from item reactivation. Error bars = SEM across subjects, +*p* < .1, \**p* < .05, \*\**p* < .01, \*\*\**p* < .001 (not corrected for multiple comparisons).

## REFERENCES

Binder JR, Desai RH. 2011. The neurobiology of semantic memory. Trends Cogn Sci. 15:527–536.

Bird CM, Keidel JL, Ing LP, Horner AJ, Burgess N. 2015. Consolidation of complex events via reinstatement in posterior cingulate cortex. J Neurosci. 35:14426–14434.

Bonnici HM, Richter FR, Yazar Y, Simons JS. 2016. Multimodal feature integration in the angular gyrus during episodic and semantic retrieval. J Neurosci. 36:5462–5471.

Bouwmeester S, Verkoeijen PPJL. 2011. Why do some children benefit more from testing than others? Gist trace processing to explain the testing effect. J Mem Lang. 65:32–41.

Brainard DH. 1997. The psychophysics toolbox. Spat Vis. 10:433–436.

Brainerd CJ, Reyna VF. 1996. Mere memory testing creates false memories in children. Dev Psychol. 32:467–478.

Bridge DJ, Paller KA. 2012. Neural correlates of reactivation and retrieval-induced distortion. J Neurosci. 32:12144–12151.

Buckner RL, Wheeler ME. 2001. The cognitive neuroscience of remembering. Nat Rev Neurosci. 2:624–634.

Chen J, Leong YC, Honey CJ, Yong CH, Norman KA, Hasson U. 2017. Shared memories reveal shared structure in neural activity across individuals. Nat Neurosci. 20:115–125.

Danker JF, Anderson JR. 2010. The ghosts of brain states past: remembering reactivates the brain regions engaged during encoding. Psychol Bull. 136:87–102.

Destrieux C, Fischl B, Dale A, Halgren E. 2010. Automatic parcellation of human cortical gyri and sulci using standard anatomical nomenclature. Neuroimage. 53:1–15.

Devereux BJ, Clarke A, Marouchos A, Tyler LK. 2013. Representational similarity analysis reveals commonalities and differences in the semantic processing of words and objects. J Neurosci. 33:18906–18916.

Fan RE, Chang KW, Hsieh CJ, Wang XR, Lin CJ. 2008. Liblinear: a library for large linear classification. J Mach Learn Res. 9:1871–1874.

Guerin SA, Robbins CA, Gilmore AW, Schacter DL. 2012. Interactions between visual attention and episodic retrieval: dissociable contributions of parietal regions during gist-based false recognition. Neuron. 75:1122–1134.

Hutchinson JB, Uncapher MR, Wagner AD. 2009. Posterior parietal cortex and episodic retrieval: convergent and divergent effects of attention and memory. Learn Mem. 16:343–356.

Ibos G, Freedman DJ. 2014. Dynamic integration of task-relevant visual features in posterior parietal cortex. Neuron. 83:1468–1480.

Karpicke JD, Roediger HL. 2008. The critical importance of retrieval for learning. Science. 319:966–968.

Koen JD, Rugg MD. 2016. Memory reactivation predicts resistance to retroactive interference: evidence from multivariate classification and pattern similarity analyses. J Neurosci. 36:4389–4399.

Kuhl BA, Chun MM. 2014. Successful remembering elicits event-specific activity patterns in lateral parietal cortex. J Neurosci. 34:8051–8060.

Kuhl BA, Rissman J, Chun MM, Wagner AD. 2011. Fidelity of neural reactivation reveals competition between memories. Proc Natl Acad Sci. 108:5903–5908.

Kuhl BA, Johnson MK, Chun MM. 2013. Dissociable neural mechanisms for goal-directed versus incidental memory reactivation. J Neurosci. 33:16099–16109.

Lee H, Kuhl BA. 2016. Reconstructing perceived and retrieved faces from activity patterns in lateral parietal cortex. J Neurosci. 36:6069–6082.

Liu XL, Liang P, Li K, Reder LM. 2014. Uncovering the neural mechanisms underlying learning from tests. PLoS One. 9:e92025.

Mack ML, Preston AR. 2016. Decisions about the past are guided by reinstatement of specific memories in the hippocampus and perirhinal cortex. Neuroimage. 127:144–157.

McDermott KB. 2006. Paradoxical effects of testing: repeated retrieval attempts enhance the likelihood of later accurate and false recall. Mem Cognit. 34:261–267.

Payne DG, Elie CJ, Blackwell JM, Neuschatz JS. 1996. Memory illusions: recalling recognizing, and recollecting events that never occurred. J Mem Lang. 35:261–285.

Polyn SM, Natu VS, Cohen JD, Norman KA. 2005. Category-specific cortical activity precedes retrieval during memory search. Science. 310:1963–1966.

Richter FR, Cooper RA, Bays PM, Simons JS. 2016. Distinct neural mechanisms underlie the success, precision, and vividness of episodic memory. ELife. 5:e18260.

Rissman J, Wagner AD. 2012. Distributed representations in memory: insights from functional brain imaging. Annu Rev Psychol. 63:101–128.

Roediger HL, Butler AC. 2011. The critical role of retrieval practice in long-term retention. Trends Cogn Sci. 15:20–27.

Roediger HL, McDermott KB. 1995. Creating false memories: remembering words not presented in lists. J Exp Psychol Learn Mem Cogn. 21:803–814.

Rugg MD, King DR. 2017. Ventral lateral parietal cortex and episodic memory retrieval. Cortex. doi:10.1016/j.cortex.2017.07.012.

Schlichting ML, Preston AR. 2014. Memory reactivation during rest supports upcoming learning of related content. Proc Natl Acad Sci. 111:15845–15850.

Shimamura AP. 2011. Episodic retrieval and the cortical binding of relational activity. Cogn Affect Behav Neurosci. 11:277–291.

St. Jacques PL, Olm C, Schacter DL. 2013. Neural mechanisms of reactivation-induced updating that enhance and distort memory. Proc Natl Acad Sci. 110:19671–19678.

St-Laurent M, Abdi H, Buchsbaum BR. 2015. Distributed patterns of reactivation predict vividness of recollection. J Cogn Neurosci. 27: 2000–2018.

Thakral PP, Wang TH, Rugg MD. 2017. Decoding the content of recollection within the core recollection network and beyond. Cortex. 91:101–113.

Tomita H, Ohbayashi M, Nakahara K, Hasegawa I, Miyashita Y. 1999. Top-down signal from prefrontal cortex in executive control of memory retrieval. Nature. 401: 699–703.

van den Broek GSE, Takashima A, Segers E, Fernández G, Verhoeven L. 2013. Neural correlates of testing effects in vocabulary learning. Neuroimage. 78:94–102.

van den Broek GSE, Takashima A, Wiklund-Hörnqvist C, Wirebring LK, Segers E, Verhoeven L, Nyberg L. 2016. Neurocognitive mechanisms of the testing effect: a review. Trends Neurosci Educ. 5:52–66.

Verkoeijen PPJL, Bouwmeester S, Camp G. 2012. A short-term testing effect in cross-language recognition. Psychol Sci. 23:567–571.

Wagner IC, van Buuren M, Kroes MCW, Gutteling TP, van der Linden M, Morris RG, Fernández G. 2015. Schematic memory components converge within angular gyrus during retrieval. ELife. 4:e09668.

Wheeler ME, Petersen SE, Buckner RL. 2000. Memory’s echo: vivid remembering reactivates sensory-specific cortex. Proc Natl Acad Sci. 97:11125–11129.

Wimber M, Alink A, Charest I, Kriegeskorte N, Anderson MC. 2015. Retrieval induces adaptive forgetting of competing memories via cortical pattern suppression. Nat Neurosci. 18:582–589.

Wing EA, Ritchey M, Cabeza R. 2015. Reinstatement of individual past events revealed by the similarity of distributed activation patterns during encoding and retrieval. J Cogn Neurosci. 27:679–691.

Wirebring LK, Wiklund-Hornqvist C, Eriksson J, Andersson M, Jonsson B, Nyberg L. 2015. Lesser neural pattern similarity across repeated tests is associated with better long-term memory retention. J Neurosci. 35:9595–9602.

Xiao X, Dong Q, Gao J, Men W, Poldrack RA, Xue G. 2017. Transformed neural pattern reinstatement during episodic memory retrieval. J Neurosci. 37:2986–2998.

Yeo BTT, Krienen FM, Sepulcre J, Sabuncu MR, Lashkari D, Hollinshead M, Roffman JL, Smoller JW, Zöllei L, Polimeni JR, et al. 2011. The organization of the human cerebral cortex estimated by intrinsic functional connectivity. J Neurophysiol. 106:1125–1165.

